# Potent Cas9 inhibition in bacterial and human cells by new anti-CRISPR protein families

**DOI:** 10.1101/350504

**Authors:** Jooyoung Lee, Aamir Mir, Alireza Edraki, Bianca Garcia, Nadia Amrani, Hannah E. Lou, Ildar Gainetdinov, April Pawluk, Raed Ibraheim, Xin D. Gao, Pengpeng Liu, Alan R. Davidson, Karen L. Maxwell, Erik J. Sontheimer

## Abstract

CRISPR-Cas systems are widely used for genome engineering technologies, and in their natural setting, they play crucial roles in bacterial and archaeal adaptive immunity, protecting against phages and other mobile genetic elements. Previously we discovered bacteriophage-encoded Cas9-specific anti-CRISPR (Acr) proteins that serve as countermeasures against host bacterial immunity by inactivating their CRISPR-Cas systems^1^. We hypothesized that the evolutionary advantages conferred by anti-CRISPRs would drive the widespread occurrence of these proteins in nature^2–4^. We have identified new anti-CRISPRs using the bioinformatic approach that successfully identified previous Acr proteins^1^ against *Neisseria meningitidis* Cas9 (NmeCas9). In this work we report two novel anti-CRISPR families in strains of *Haemophilus parainfluenzae* and *Simonsiella muelleri*, both of which harbor type II-C CRISPR-Cas systems^5^. We characterize the type II-C Cas9 orthologs from *H. parainfluenzae* and *S. muelleri*, show that the newly identified Acrs are able to inhibit these systems, and define important features of their inhibitory mechanisms. The *S. muelleri* Acr is the most potent NmeCas9 inhibitor identified to date. Although inhibition of NmeCas9 by anti-CRISPRs from *H. parainfluenzae* and *S. muelleri* reveals cross-species inhibitory activity, more distantly related type II-C Cas9s are not inhibited by these proteins. The specificities of anti-CRISPRs and divergent Cas9s appear to reflect co-evolution of their strategies to combat or evade each other. Finally, we validate these new anti-CRISPR proteins as potent off-switches for Cas9 genome engineering applications.

---

Clustered, regularly interspaced, short, palindromic repeats (CRISPR) and their CRISPR-associated (*cas*) genes constitute a prokaryotic adaptive immune defense system against foreign genetic elements such as phages and plasmids^6–8^. The components of CRISPR-Cas systems that allow recognition and destruction of invading genetic elements are extremely diverse, and form the basis for the current CRISPR-Cas classification framework^9^, which includes two broad classes, six types, and multiple subtypes. In class 1 CRISPR systems, effector modules form a multi-protein complex, whereas class 2 systems use a single effector protein to target foreign nucleic acids. Cas9 is an effector protein in the best-studied class 2 system, type II, which is further divided into three subtypes (II-A, -B, and -C) based on Cas9 phylogeny and the presence or absence of additional adaptation-related Cas proteins^9^. Cas9 is a single-component, RNA-guided endonuclease that employs the CRISPR RNA (crRNA) as a sequence-specific guide to target foreign DNA^10^, with the help of a trans-activating RNA (tracrRNA)^11^, which can be fused to the crRNA to form a single guide RNA (sgRNA)^12^. The robustness and ease of Cas9 programmability have greatly facilitated its rapid adoption in genome editing and modulation^13^.

Although Cas9s have attracted unprecedented attention for genome engineering applications, their natural function in bacterial defense plays a crucial role in the ongoing battle against phages and other invading mobile genetic elements (MGEs). As countermeasures against such a powerful barrier, phages and MGEs have evolved numerous, distinct strategies to overcome bacterial defenses^14^. Anti-CRISPR (Acr) proteins provide one way to directly disarm CRISPR-Cas systems. The existence of Acrs was first shown in phages that successfully infect *Pseudomonas aeruginosa* strains despite the presence of active type I CRISPR-Cas systems and phage-matched CRISPR spacers^15^. Type I Acr families do not share common structural similarities or sequences, but are frequently encoded adjacent to putative transcriptional regulator genes known as anti-CRISPR-associated (*aca*) genes^16^. The first type II-specific *acr* genes were identified as previously uncharacterized open reading frames (ORFs) adjacent to predicted *aca* genes in MGEs of bacteria harbouring type II CRISPR-Cas systems^1^. Additional Acrs have been found by identifying candidate *acr* genes in lysogens embedded within genomes harboring potentially self-targeting type II CRISPR systems^17^, or by screening lytic phages for the ability to resist type II CRISPR defenses^18^. Type II Acrs are of particular interest because they can potentially provide temporal, spatial, or conditional control over Cas9-based applications.

Thus far, three families of type II-C Acrs^1^ and five families of type II-A Acrs^17, 18^ have been reported, and inhibitory mechanisms are known in a few cases^1, 17, 19^. For instance, AcrIIA4_*Lmo*_, a type II-A Acr that can inhibit the most widely-used Cas9 ortholog from *Streptococcus pyogenes* (SpyCas9), prevents Cas9 DNA binding^17^ by occupying the protospacer adjacent motif (PAM)-interacting domain and masking the RuvC nuclease domain, in part via DNA mimicry^20–22^. Conversely, a type II-C Acr, AcrIIC1_*𝒩me*_, does not prevent target DNA binding by *𝒩eisseria meningitidis* Cas9 (NmeCas9, from strain 8013), but rather binds and inhibits the enzyme’s HNH nuclease domain^19^. Yet another type II-C Cas9 inhibitor, AcrIIC3_*𝒩me*_, prevents target DNA binding1 in a manner that is accompanied by NmeCas9 dimerization^19^.

Because Acrs provide obvious fitness advantages^23^ to phages and MGEs, we hypothesized that many more type II Acrs likely remain to be discovered. Here, we identify two new type II-C Cas9 inhibitors from strains of *H. parainfluenzae* (AcrIIC4_*Hpa*_) and *S. muelleri* (AcrIIC5_*Smu*_). We characterize their cognate Cas9 proteins from *H. parainfluenzae* and *S. muelleri* and show that these proteins are functional *in vivo* and *in vitro*. Further, we show that AcrIIC5_*Smu*_ is the most potent NmeCas9 inhibitor reported to date. While both of these Acrs inhibit DNA binding by Cas9, including during mammalian genome editing applications, they differ in their phylogenetic ranges of Cas9 inhibition.

## Results

### Identification of novel type II-C anti-CRISPR proteins

We previously developed a ‘guilt-by-association’ bioinformatics approach that allowed the identification of novel families of anti-CRISPR proteins encoded in phages and MGEs of diverse bacterial species^1, 16^. In this pipeline, new *acr* gene candidates are identified by their proximity to predicted helix-turn-helix (HTH) transcriptional regulator genes known as anti-CRISPR associated (*aca*) genes^16^. We identified ORFs encoding uncharacterized small [~50-150 amino acids (aa)] proteins immediately upstream of *aca* homologues, focusing on genomic regions near putative phage- or MGE-associated sequences^3, 4, 24^. These criteria led us to focus on two putative Acr candidates: an 88 aa hypothetical protein in the genome of *H. parainfluenzae* strain 146_HPAR (NCBI RefSeq accession WP_049372635) and a 120aa hypothetical protein in the genome of *S. muelleri* strain ATCC 29453 (WP_002642161.1; Supplementary Table 1). Both are located upstream of apparent *aca* orthologs, near potential phage or MGE genes (Fig. 1a), and both strains encode predicted type II-C CRISPR-Cas machineries with Cas9 orthologs that exhibit 59% and 62% identity with NmeCas9, respectively (Supplementary Table 2). Based on these similarities, the previously established abilities of some type II anti-CRISPRs to inhibit Cas9 orthologs outside of their host species^1, 17, 19, 25^, and the existence of apparent orthologues of the *S. muelleri* candidate Acr in multiple examples from *𝒩eisseria* (Supplementary Table 3), we first tested for anti-CRISPR activity against the well-characterized NmeCas9. We cloned each candidate Acr sequence into a bacterial expression vector, purified recombinant proteins from *Escherichia coli*, and tested their abilities to prevent DNA cleavage by NmeCas9 *in vitro* (Fig. 1b). When each of the purified candidate Acrs was added to parallel reactions, cleavage was inhibited in a concentration-dependent manner, with complete inhibition being reached at ~15-fold (*H. parainfluenzae* candidate) and ~4-fold (*S. muelleri* candidate) molar excess of Acr (Fig. 1b). Incubation with AcrE2, an 84aa type I-E anti-CRISPR^1, 16^ included as a negative control, did not affect target DNA cleavage by NmeCas9. When we compared the ability of Acrs to inhibit DNA cleavage when first added to the apo or sgRNA-loaded forms of NmeCas9, both candidate Acrs inhibited the two forms of NmeCas9 to a comparable extent (Supplementary Fig. 1). This observation is in contrast to previously described orthologous anti-CRISPRs AcrIIC1_*Boe*_ and AcrIIC1_*𝒩me*_, which were less potent when added to the NmeCas9:sgRNA complex (Supplementary Fig. 1). Because these initial tests confirmed the anti-CRISPR activities of the two candidates from *H. parainfluenzae* and *S. muelleri*, we named them AcrIIC4_*Hpa*_ and AcrIIC5_*Smu*_, respectively, to conform with established Acr nomenclature^4, 26^.

**Figure 1.**
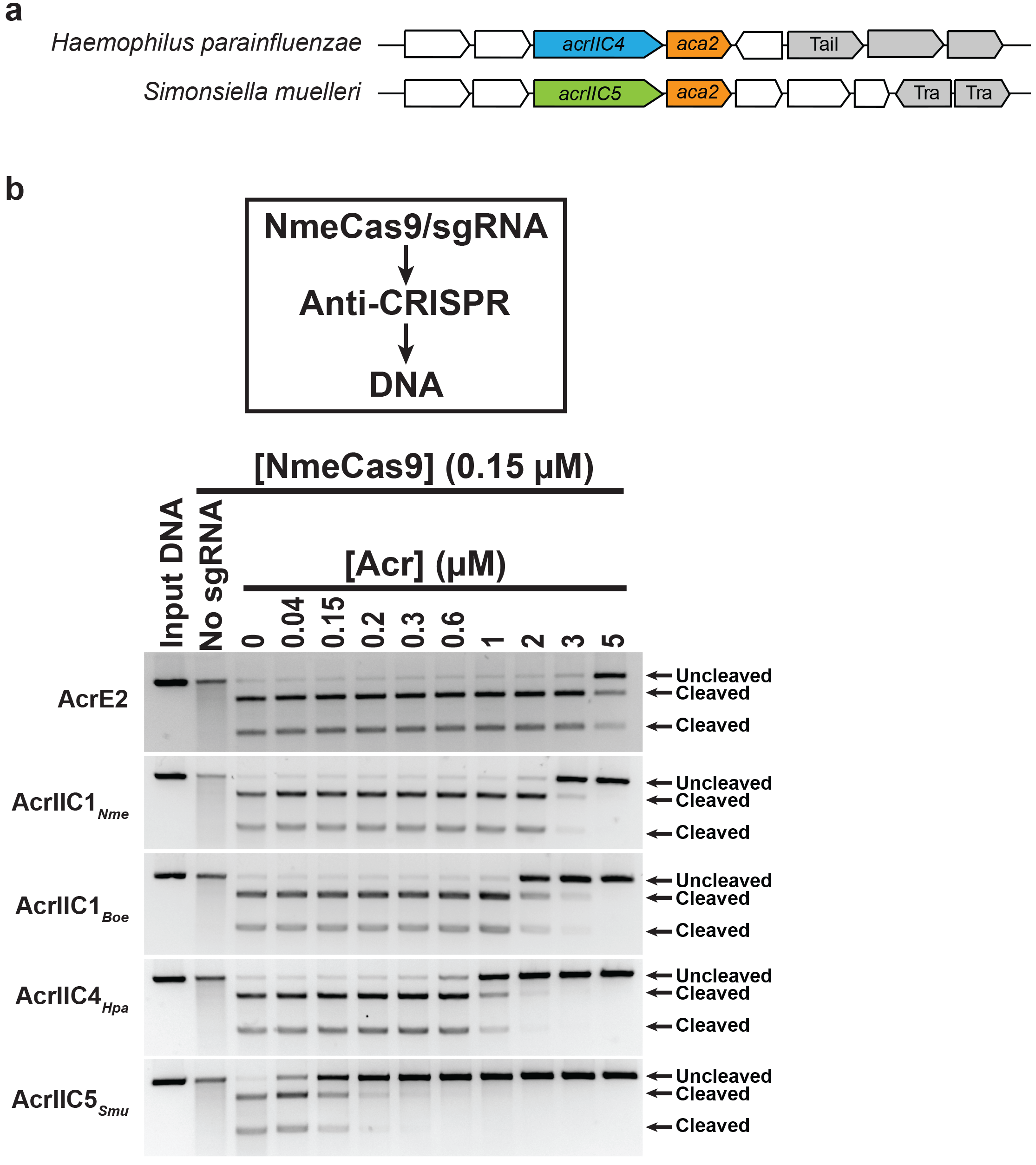
Identification and *in vitro* validation of two anti-CRISPR protein families. **a,** Schematic of candidate anti-CRISPR proteins and *aca* genes in the genomic context of *H. parainfluenzae* (AcrIIC4_*Hpa*_) and *S. muelleri (*AcrIIC5*Smu*). Gray-colored genes are associated with mobile DNA, and known gene functions are annotated as following: ‘Tail’ is involved in phage tail morphogenesis; ‘Tra’ is a transposase. Arrows are not drawn to scale. **b,** *In vitro* cleavage of target DNA by the NmeCas9-sgRNA complex in the presence of anti-CRISPR protein. A linearized plasmid with a protospacer and PAM sequence was subjected to *in vitro* digestion by purified, recombinant, sgRNA-programmed NmeCas9. Preformed NmeCas9-sgRNA RNP complex was incubated with purified anti-CRISPR proteins as indicated with AcrE2 as a negative control, AcrIIC1 as a positive control, and candidate Acrs. Molarities of anti-CRISPR protein (relative to constant Cas9 molarity) are shown at the top of each lane, and mobilities of input and cleaved DNAs are denoted on the right.

### *H. parainfluenzae* and *S. muelleri* encode type II-C CRISPR-Cas systems that function *in vitro*

Anti-CRISPRs are most likely to inhibit the Cas9 ortholog expressed by the same strain, but to our knowledge, little was known about the Cas9 orthologs from *H. parainfluenzae* and *S. muelleri*. To address this, we characterized these type II-C CRISPR-Cas systems (Fig. 2a). First, we identified a 1,054aa *cas9* ORF in *H. parainfluenzae* DSM 8978, a strain closely related to *H. parainfluenzae* 146_HPAR for which genomic DNA was available. We identified a predicted tracrRNA adjacent to *cas9*, and noted that the CRISPR repeat sequence included a likely minimal promoter that initiates transcription in the flanking spacer, as found previously with other type II-C systems^5^ (Supplementary Fig. 2a). The predicted transcriptional start site would yield a crRNA with a 24-nt spacer, similar to *𝒩. meningitidis* strain 8013^27^. We then used tracrRNA/crRNA complementarity to predict an sgRNA scaffold. These components were then used to generate expression constructs for recombinant HpaCas9 in *E. coli*, and for its sgRNA via in *vitro* transcription, for biochemical analyses (below).

**Figure 2.**
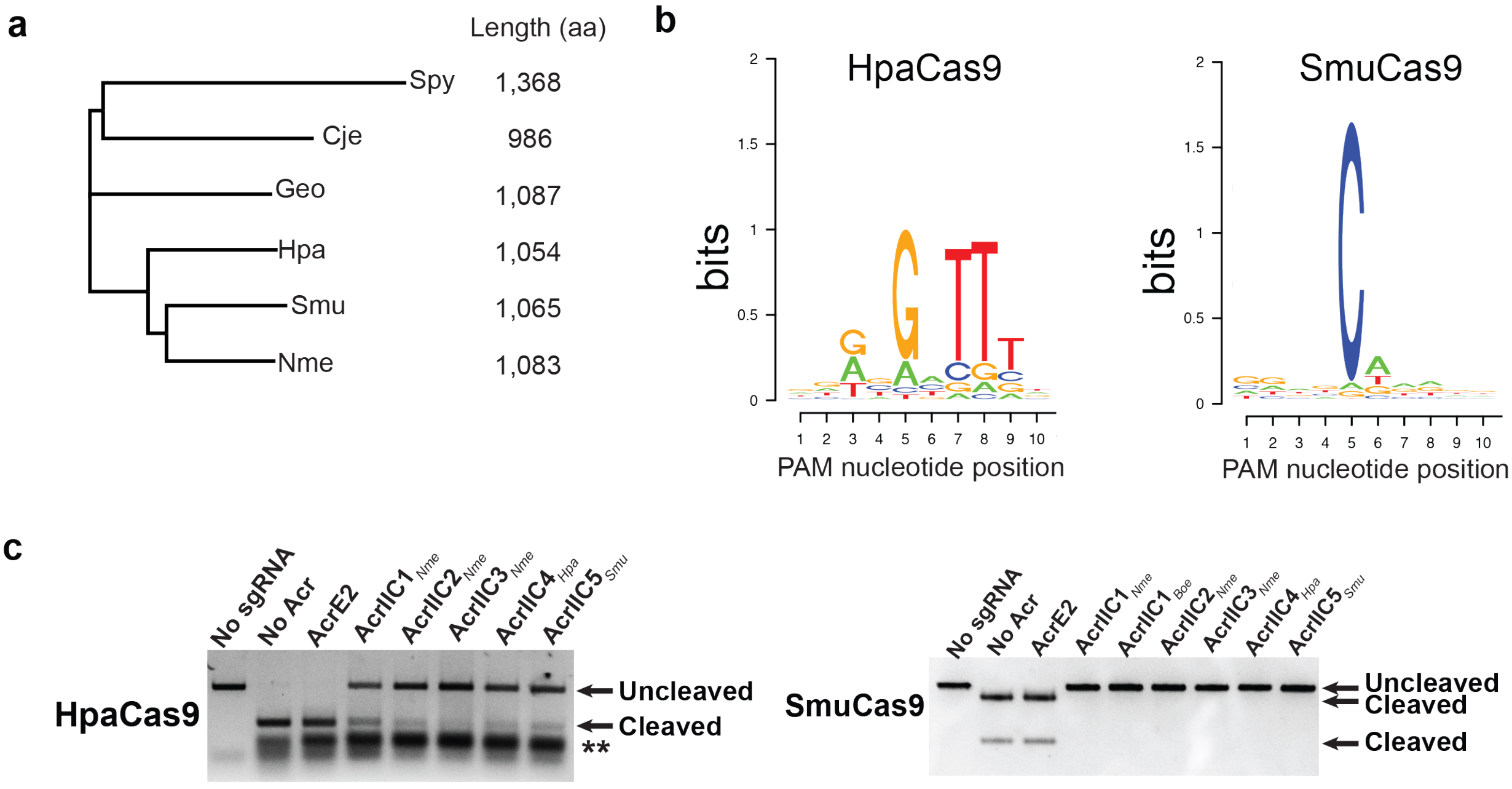
A diversity of Cas9 orthologs and breadth of inhibition by anti-CRISPR proteins. **a,** A phylogenetic tree of type II-C Cas9 orthologs from *𝒩. meningitidis, C. jejuni, G. stearothermophilus, H. parainfluenzae,* and *S. muelleri.* Cas9 from *S. pyogenes* is used as an outgroup. **b,** PAM preferences for *H. parainfluenzae* (left) and *S. muelleri* (right) Cas9 proteins. Frequencies of nucleotides at each PAM position were calculated and plotted as a WebLogo. **c,** Validation of HpaCas9 and SmuCas9 cleavage activity and inhibition by anti-CRISPR proteins *in vitro*. The double asterisk denotes sgRNA.

Unlike *H. parainfluenzae* DSM 8978, the CRISPR-*cas* locus of *S. muelleri* ATCC 29453 appeared to be degenerate (Supplementary Fig. 2a). There is no apparent *cas1* ORF, and the *cas2* ORF lacks a canonical ATG start codon. However, the *cas9* ORF (1,065aa) is intact and has all the predicted functional domains found in other Cas9 orthologs, which suggested that SmuCas9 itself might be active. When we attempted to define an appropriate guide RNA scaffold for SmuCas9, we could not predict its tracrRNA (based in part on crRNA complementarity) from nearby genomic sequences. Instead, we found an IS5 integrase upstream of *cas9*, where a *tracrRNA* locus is often observed (Supplementary Fig. 2a). Although we sequenced ~2 kb flanking the CRISPR locus to fill gaps in the genome assembly, we could not detect a tracrRNA sequence. As an alternative, we took advantage of the non-orthogonality of sgRNAs from closely related Cas9 orthologs^28^, ^29^ and used the NmeCas9 sgRNA to test the cleavage activity of SmuCas9.

To define the PAM requirements for HpaCas9 and SmuCas9, a library of short DNA fragments containing a unique protospacer and 10-nt randomized PAM sequences was subjected to *in vitro* digestion using purified, recombinant Cas9 proteins and T7-transcribed sgRNAs. Next, digested products were gel-purified and deep-sequenced. PAM sequences were identified from the resulting sequencing data based on the frequency of nucleotides at each position of the digested products. HpaCas9 had strong preference for 5’-N_4_GATT-3’ PAM sequence (Fig. 2b). Notably, this PAM is similar to the consensus PAM sequence of NmeCas9^27, 28, 30–32^. We extracted spacer sequences from the *H. parainfluenzae* 146_HPAR CRISPR locus and identified two candidate protospacers. When we aligned the nucleotide sequences adjacent to the protospacers, we noted that both contained a 5’-N_4_GATT-3’, which is consistent with the PAM discovered *in vitro* (Fig. 2b and Supplementary Fig. 2b). We found that SmuCas9 had strong preference for the 5’-N_4_C-3’ PAM sequence (Fig. 2b). This single cytosine at the 5^th^ position from the protospacer appears to be the most critical PAM nucleotide by far, although moderate preferences for other nucleotides at other positions cannot be excluded from this analysis. We validated these putative PAMs by performing *in vitro* cleavage of a non-degenerate substrate and confirmed efficient cleavage of a DNA target bearing a 5’-N_4_GATT-3’ PAM for HpaCas9 and a 5’-N_4_C-3’ PAM for SmuCas9 (Fig. 2c).

### AcrIIC4_*Hpa*_ and AcrIIC5_*Smu*_ inhibit their native, cognate Cas9 proteins and close orthologs *in vitro* and in bacteria

We next examined the ability of AcrIIC4_*Hpa*_ and AcrIIC5_*Smu*_ to inhibit HpaCas9 and SmuCas9, which share 59% and 62% sequence identity with NmeCas9, respectively. Our *in vitro* DNA cleavage analyses show that these Acrs can inactivate their cognate Cas9 proteins (Fig. 2c). Given that some type II Acrs can inhibit orthologous Cas9 within the same subtype^17, 18, 33^, we tested *𝒩eisseria* representatives of the three other type II-C Acr families (AcrIIC1_*𝒩me*_, AcrIIC2_*𝒩me*_, and AcrIIC3_*𝒩me*_) for inhibition of these two newly characterized Cas9 proteins. We found that all three of these previously characterized Acrs inhibit the DNA cleavage activity of both HpaCas9 and SmuCas9 (Fig. 2c).

To further characterize the physical interactions of AcrIIC4_*Hpa*_ and AcrIIC5_*Smu*_ with HpaCas9 and SmuCas9, we co-expressed each 6xHis-tagged Cas9 together with untagged Acr proteins in *E. coli*. Using Ni^2+^-affinity chromatography, we determined that AcrIIC4_*Hpa*_ directly bound HpaCas9 and SmuCas9 (Fig. 3a). This is similar to the results observed for the previously characterized type II-C Acrs, which are known to bind to NmeCas9^19, 33^. By contrast, AcrIIC5_*Smu*_ did not co-purify with any of the tested Cas9 proteins under these conditions (Fig. 3a).

**Figure 3.**
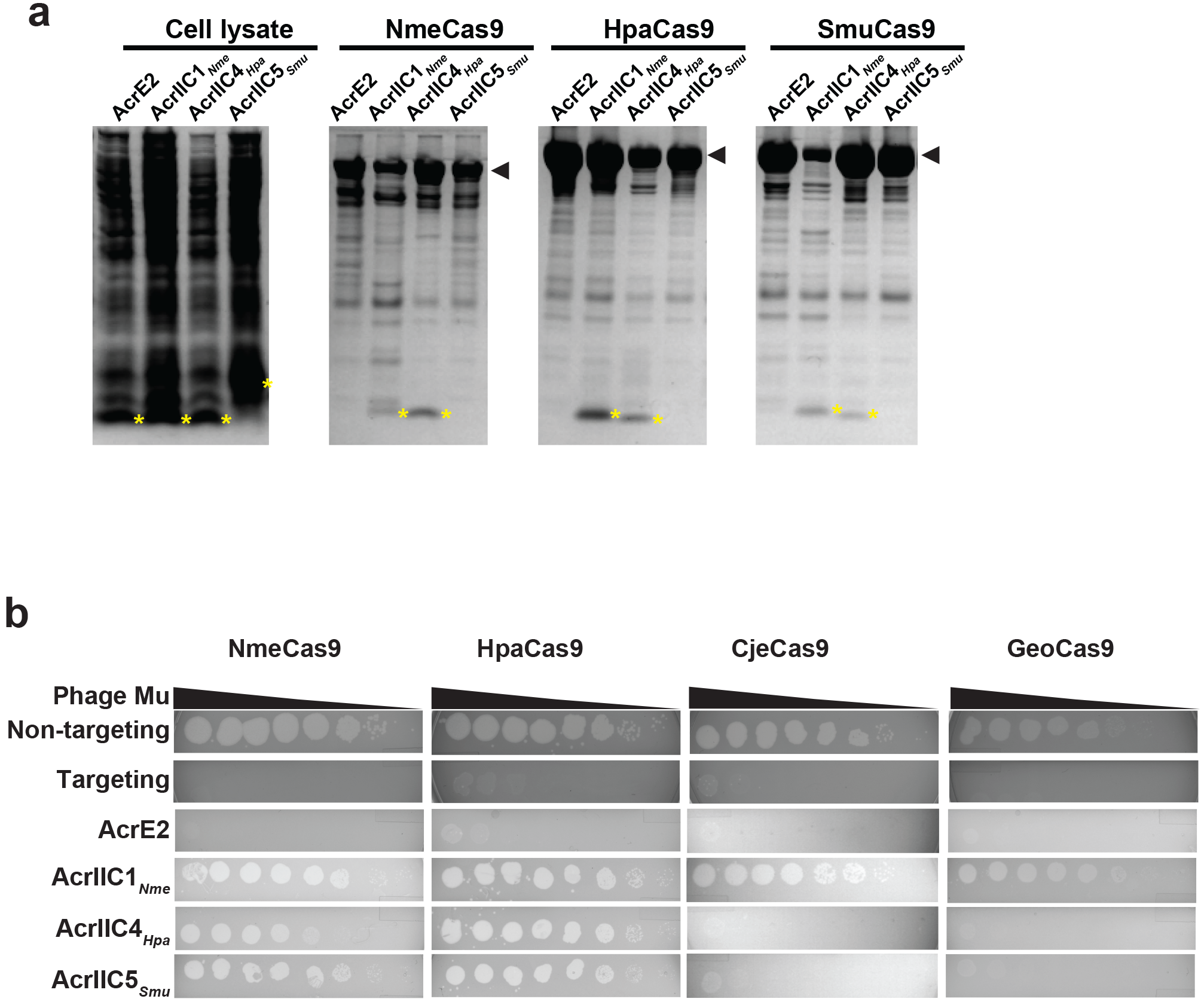
Plaquing of *E. coli* phage Mu targeted by the Nme, Hpa, Cje, or Geo Cas9 in the presence of the anti-CRISPR proteins. **a,** Interaction between Acrs and NmeCas9, HpaCas9, and SmuCas9 is visualized by Coomassie staining after co-purification of each 6xHis-tagged Cas9 and untagged Acrs from *E. coli*. Each Cas9 ortholog and anti-CRISPRs are indicated as arrowhead and yellow asterisks, respectively. **b,** Ten-fold serial dilutions of phage Mu lysate were spotted on lawns of bacteria expressing the indicated Acr proteins. Data are from one plate representative of ≥3 replicates.

Previous work has shown that some Acrs, such as AcrIIC1 family members, inhibit Cas9s from *Campylobacter jejuni* (CjeCas9) and *Geobacillus stearothermophilus* (GeoCas9), in addition to NmeCas9^34^. CjeCas9 shares 32% sequence identity with NmeCas9, and GeoCas9 shares 39% (Supplementary Table 2). To determine the range of activity of AcrIIC4_*Hpa*_ and AcrIIC5_*Smu*_, we tested their inhibitory effects on type II-C Cas9s that have been validated for mammalian genome editing (Supplementary Fig. 2c). Despite the abilities of both AcrIIC4_*Hpa*_ and AcrIIC5_*Smu*_ to inhibit DNA cleavage by NmeCas9 *in vitro*, neither prevented target DNA cleavage by CjeCas9 or GeoCas9.

To confirm these *in vitro* results, we also performed *E. coli*-based phage targeting assays to assess the ability of AcrIIC4_*Hpa*_ and AcrIIC5_*Smu*_ to inhibit the activity of the various Cas9 orthologs. In this assay, Cas9 expressed from a plasmid in *E. coli* with an sgRNA that targets phage Mu led to a reduction in phage titer of ~106 plaque-forming units (pfu)/mL (Fig. 3b). AcrIIC4_*Hpa*_ expression completely inhibited the activity of HpaCas9 and decreased the activity of NmeCas9 by ~100-fold (Fig. 3b and Supplementary Fig. 3). Similarly, AcrIIC5_*Smu*_ completely inhibited the activity of both NmeCas9 and HpaCas9 (Fig. 3b), allowing phage Mu to plaque robustly. We were unable to test inhibition of SmuCas9 activity in *E. coli* because it failed to interfere with phage Mu plaquing even in the absence of Acr proteins, perhaps due to compromised function *in vivo* with the non-cognate NmeCas9 sgRNAs. Consistent with the *in vitro* results, neither AcrIIC4_*Hpa*_ nor AcrIIC5_*Smu*_ inhibited phage targeting by GeoCas9 or CjeCas9 (Fig. 3b). These results, together with the *in vitro* DNA cleavage assays (Fig. 1), indicate that AcrIIC4_*Hpa*_ and AcrIIC5_*Smu*_ exhibit cross-species inhibitor activity (based on NmeCas9 inhibition) but have a narrower inhibitory spectrum than AcrIIC1^19^.

### AcrIIC4_*Hpa*_ and AcrIIC5_*Smu*_ inhibit NmeCas9-mediated genome editing in mammalian cells

Validation of anti-CRISPR activity *in vitro* and in bacteria prompted us to test whether AcrIIC4_*Hpa*_ and AcrIIC5_*Smu*_ inhibit genome editing in mammalian cells. First, we used co-immunoprecipitation experiments to confirm that the NmeCas9/AcrIIC4_*Hpa*_ physical interaction observed with recombinant proteins *in vitro* (Fig. 3a) can also be detected in lysates from mammalian cells (Supplementary Fig. 4a). Consistent with AcrIIC5_*Smu*_ inhibition of NmeCas9 DNA cleavage activity *in vitro* (Fig. 1b), we also detected AcrIIC5_*Smu*_/NmeCas9 co-immunoprecipitation in mammalian lysates (Supplementary Fig. 4a), even though purified, recombinant NmeCas9 did not pull down recombinant AcrIIC5_*Smu*_ expressed in *E. coli* (Fig. 3a). To assess the inhibition of NmeCas9 genome editing, we co-transfected HEK293T cells transiently with plasmids expressing anti-CRISPR protein, NmeCas9 and sgRNAs targeting genomic sites. We then used T7 endonuclease 1 (T7E1) digestion to estimate genome editing efficiency. In agreement with our *in vitro* data, expression of AcrIIC4_*Hpa*_ or AcrIIC5_*Smu*_ reduced NmeCas9-mediated mutagenesis to undetectable levels at both tested sites (Fig. 4a). In contrast, they had no effect on genome editing at the same genomic sites by SpyCas9, which belongs to the type II-A CRISPR-Cas subtype and is very distantly related to NmeCas9 (Fig. 4a). Titration of plasmids expressing AcrIIC4_*Hpa*_ or AcrIIC5_*Smu*_ demonstrated potency against NmeCas9 that was comparable or superior to that of AcrIIC3_*𝒩me*_ (Supplementary Fig. 4b), which had previously been defined as the most potent NmeCas9 inhibitor in mammalian cells^1^. For more rigorous quantitation of NmeCas9 editing, we used targeted deep sequencing at a distinct editing site (NTS1C) and detected little to no editing at higher doses of AcrIIC4_*Hpa*_ or AcrIIC5_*Smu*_ plasmids (Supplementary Fig. 4c).

**Figure 4.**
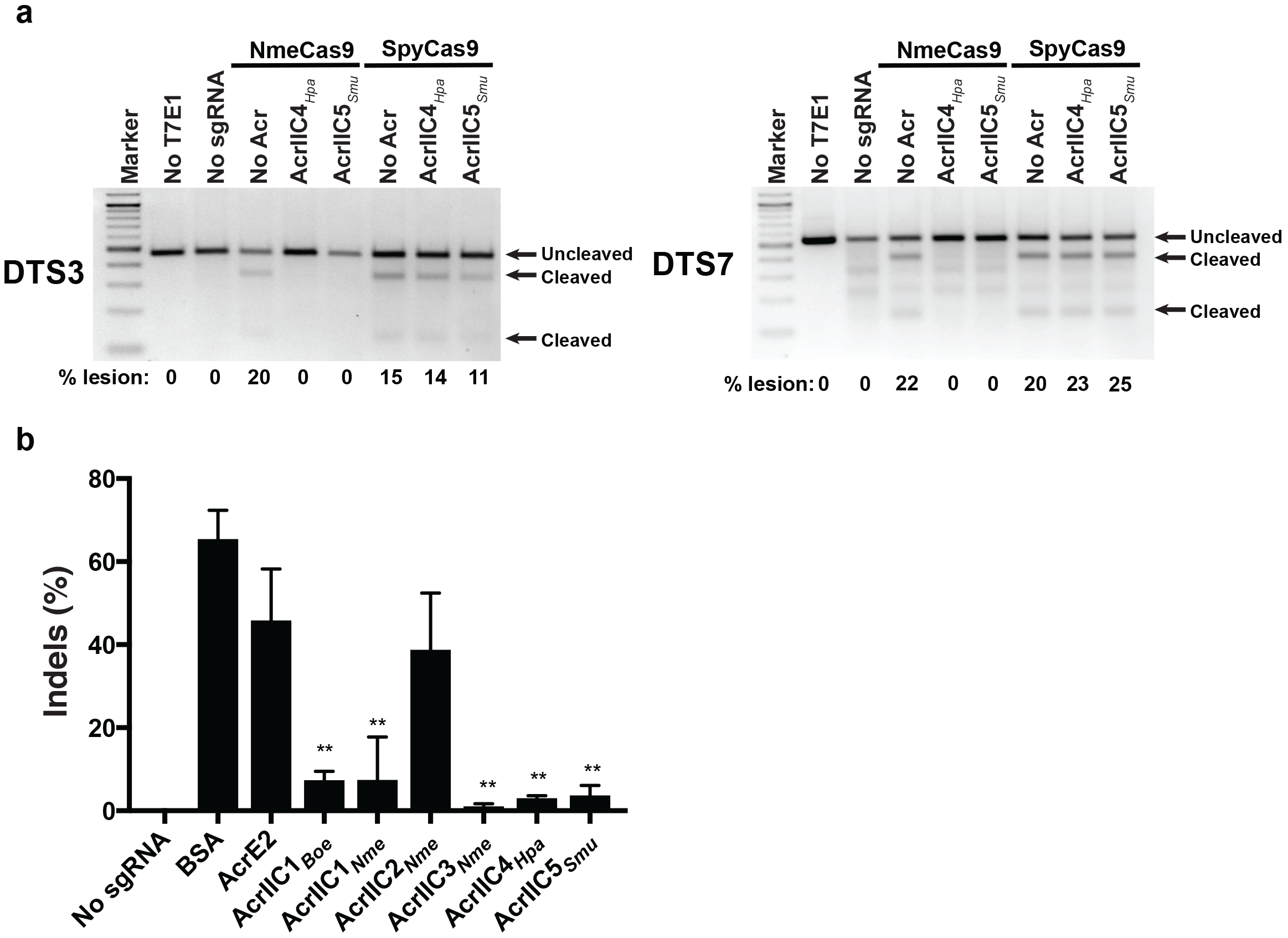
AcrIIC4_*Hpa*_ and AcrIIC5_*Smu*_ inhibit genome editing by NmeCas9 in human cells. **a,** T7E1 assays of NmeCas9 or SpyCas9 editing efficiencies at two dual target sites (DTS3 and DTS7) upon transient plasmid transfection of human HEK293T cells. Constructs encoding anti-CRISPR proteins were co-transfected as indicated at the top of each lane. Mobilities of edited and unedited bands are indicated to the right and editing efficiencies (“% lesion”) are given at the bottom of each lane. The figure is a representative of three replicates. **b,** A bar graph of editing efficiencies measured by TIDE analysis upon RNP delivery of NmeCas9-sgRNA and Acr into HEK293T cells. Statistical significance was determined by two-tailed paired student’s t-test. Mean and standard deviations of three biological replicates are indicated with lines (*; *p* < 0.05, **; *p* < 0.01, ***; *p* < 0.001).

We previously noted a discrepancy in the potency of AcrIIC3_*𝒩me*_, which was least active in inhibiting *𝒩. meningitidis* transformation but was most potent in cultured human cells^1^. To address whether anti-CRISPR expression or stability correlates with inhibitory effect in mammalian cells, we estimated Acr protein abundance by western blots using lysates from HEK293T cells transiently transfected with Acr expression plasmids (identical in all respects other than Acr ORF). Inhibition potency correlated well with the abundance of the anti-CRISPR, with AcrIIC4_*Hpa*_ and AcrIIC5_*Smu*_ showing the highest protein signal at steady state (Supplementary Fig. 4d). To bypass the difference in protein abundance, we delivered a preformed ribonucleoprotein (RNP) complex of NmeCas9, sgRNA, and each Acr to HEK293T cells by electroporation. Then, we confirmed genome editing inhibition by AcrIIC4_*Hpa*_ and AcrIIC5_*Smu*_ using tracking of indels by decomposition (TIDE) analysis35 (Fig. 4b). Acrs still displayed variations in activities even with RNP delivery, suggesting differences in protein stability, off-rate, or other intrinsic properties. Of note, however, AcrIIC4_*Hpa*_ and AcrIIC5_*Smu*_ consistently exhibited strong inhibitory potency both *in vitro* and in cultured cells (Fig. 1b and Fig. 4). Furthermore, AcrIIC4_*Hpa*_ and AcrIIC5_*Smu*_ co-expression increased the steady-state accumulation of NmeCas9 (with or without sgRNA co-expression), consistent with the possibility of a stabilizing physical interaction (Supplementary Fig. 4e). Overall, these data demonstrate that the two new anti-CRISPRs directly bind to NmeCas9 and specifically inhibit its DNA cleavage activity in human cells.

### AcrIIC4_*Hpa*_ and AcrIIC5_*Smu*_ prevent stable DNA binding by NmeCas9

Once we confirmed the anti-CRISPR inhibition of sgRNA-guided NmeCas9 DNA cleavage *in vitro* (Fig. 1) and genome editing in cells (Fig. 4), we then addressed the mechanisms of NmeCas9 inhibition by AcrIIC4_*Hpa*_ and AcrIIC5_*Smu*_. Since structural and biochemical analysis of the anti-CRISPRs characterized to date suggest diverse and unique inhibitory mechanisms^2-4, 24^, we tested multiple hypotheses: Acrs prevent sgRNA loading, DNA target binding [like AcrIIC3_*𝒩me*_^1^], or DNA target cleavage [like AcrIIC1_*𝒩me*_^1, 19^]. First, we checked whether sgRNA loading onto NmeCas9 is inhibited by either anti-CRISPR. We carried out electrophoretic mobility shift assays (EMSA) by incubating NmeCas9 and sgRNA with or without Acr, and then visualizing sgRNA mobility after native gel electrophoresis by SYBR Gold staining. In the absence of any anti-CRISPR, incubation of NmeCas9 with its cognate sgRNA resulted in a gel shift that indicates formation of a stable RNP complex (Fig. 5a). When NmeCas9 was incubated with a negative control anti-CRISPR (AcrE2) before the addition of sgRNA, NmeCas9:sgRNA complex formation was unaffected. Similarly, when incubated with AcrIIC4_*Hpa*_ and AcrIIC5_*Smu*_, efficient NmeCas9:sgRNA complex formation was again observed, suggesting that neither Acr protein significantly affected RNP assembly.

**Figure 5.**
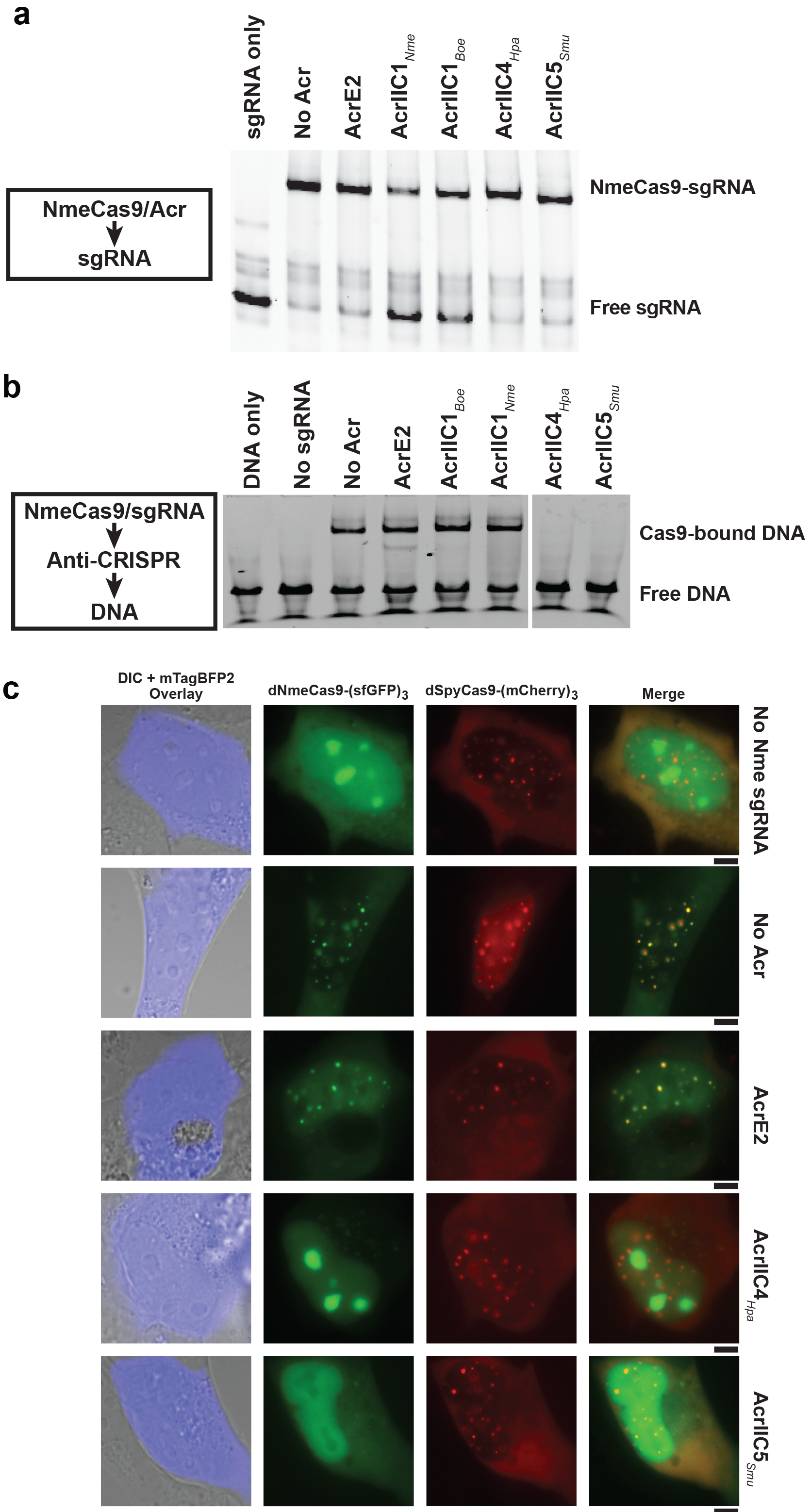
AcrIIC4_*Hpa*_ and AcrIIC5_*Smu*_ prevent stable DNA binding by NmeCas9. **a,b,** A native gel of the sgRNA visualized by SYBR gold staining (**a**) and of the FAM-labeled target DNA (**b**), both of which were added last to NmeCas9 + Acr [and in (**b**), + sgRNA] incubation. **c**, Live-cell fluorescence images of U2OS cells transiently transfected with plasmids encoding dNmeCas9-(sfGFP)_3_, dSpyCas9-(mCherry)_3_, their respective telomeric sgRNAs, and Acrs. The plasmid encoding the Acr is also marked with the blue fluorescent protein, mTagBFP2, which is overlaid on a differential interference contrast (DIC) image of each cell. The specific version of each plasmid set (with or without sgRNAs, with or without anti-CRISPRs) is given to the right of each row. First column: differential interference contrast (DIC) and mTagBFP2 imaging, overlay. Second column: dNmeCas9-(sfGFP)_3_. Third column: dSpyCas9-(mCherry)_3_. Fourth column: dNmeCas9-(sfGFP)_3_ and dSpyCas9-(mCherry)_3_, merged. Scale bars, 5 μm.

To test if target DNA engagement by the NmeCas9:sgRNA complex is prevented by either AcrIIC4_*Hpa*_ or AcrIIC5_*Smu*_, we performed EMSA and fluorescence polarization assays after incubating the RNP with each Acr, before adding target DNA (Fig. 5b and Supplementary Fig. 5a). To inhibit DNA target cleavage, we omitted divalent metal ions from the reactions. While the target DNA exhibited the expected mobility shift in the absence of Acr, or in the presence of AcrE2 or AcrIIC1_*𝒩me*_ [as expected19], both AcrIIC4_*Hpa*_ and AcrIIC5_*Smu*_ prevented NmeCas9 RNP binding to the target DNA. We also performed fluorescence polarization assays to measure the equilibrium binding constants of NmeCas9 RNP (0 - 2 μM) to target DNA (8 nM) in the presence or absence of Acrs (10 μM). As shown in Supplementary Fig. 5a, AcrIIC4_*Hpa*_ and AcrIIC5_*Smu*_ significantly impair the DNA binding activity of NmeCas9:sgRNA, confirming our EMSA results. The measured K_*d*_ of the NmeCas9 RNP to this target DNA (without Acr inhibition) is 86 ± 4 nM, similar to a previous measurement (70 ± 5 nM) with a different sgRNA/target site combination^19^. The addition of AcrIIC4_*Hpa*_ and AcrIIC5_*Smu*_ reduced apparent DNA affinity by ~9-fold (to 750 ± 150 nM) and ~6-fold (to 450 ± 50 nM), respectively (Supplementary Fig. 5a), similar to the ~10-fold inhibition of NmeCas9 DNA binding by AcrIIC3_*𝒩me*_^19^.

### AcrIIC4_*Hpa*_ and AcrIIC5_*Smu*_ are potent inhibitors of dNmeCas9-based tools in mammalian cells

Many applications (e.g. CRISPRi and CRISPRa) have been developed for catalytically inactive (“dead”) Cas9 (dCas9) derivatives fused or tethered to various effector domains^36^. To extend our findings from *in vitro* studies to mammalian cells, we tested whether AcrIIC4_*Hpa*_ and AcrIIC5_*Smu*_ prevent stable DNA binding of dNmeCas9 using previously established methods for live-cell imaging of telomeric foci. Briefly, transfection of plasmids expressing dCas9 orthologs fused to fluorescent proteins, as well as cognate sgRNAs targeting telomeres, enables telomeric foci to be visualized in U2OS cells^37^. Orthogonal dNmeCas9-(sfGFP)_3_ and dSpyCas9-(mCherry)_3_ can be used in this fashion simultaneously to bind telomeres and generate co-localizing sfGFP and mCherry telomeric foci^38^.

Transfection of a third plasmid, marked with mTagBFP2 and encoding an anti-CRISPR protein, can be used to assess the anti-CRISPR’s effects on telomeric DNA binding in live cells^1^. AcrE2 had no effect on telomeric foci formed by dNmeCas9-(sfGFP)_3_ and dSpyCas9-(mCherry)_3_, as seen previously1; however, co-expression of AcrIIC4_*Hpa*_ or AcrIIC5_*Smu*_ resulted in loss of green foci formation by dNmeCas9-(sfGFP)_3_ without abolishing the red telomeric foci formed by dSpyCas9-(mCherry)_3_ (Fig. 5c). We then quantified the number of cells exhibiting telomeric dNmeCas9-(sfGFP)_3_ foci in a blinded experimental setup (Supplementary Fig. 5b). We observed dNmeCas9-(sfGFP)_3_ foci in approximately 80% of cells in the absence of any Acr protein, in 70% of cells expressing AcrE2 protein (negative control), and in 0% of cells in the presence of AcrIIC3_*𝒩me*_ (as a positive control1, 19). We found that 0% of cells exhibited dNmeCas9-(sfGFP)_3_ telomeric foci when the two novel anti-CRISPRs were co-expressed (0 out of 78 for AcrIIC4_*Hpa*_, and 0 out of 82 for AcrIIC5_*Smu*_) (Supplementary Fig. 5b). These results confirm that AcrIIC4_*Hpa*_ and AcrIIC5_*Smu*_ inhibit stable DNA binding of dNmeCas9 in a cellular context, indicating their potential utility as potent off-switches for dNmeCas9-based applications.

As with AcrIIC4_*Hpa*_ and AcrIIC5_*Smu*_, type II anti-CRISPRs described thus far have been shown to inhibit Cas9 orthologs only within a single sub-type (II-A or II-C)^17, 18^, ^33^. Recently, SpyCas9 fusions to NmeCas9 or dNmeCas9 have been developed for the induction of precise segmental deletions, or to prevent SpyCas9 off-target editing, respectively (Bolukbasi *et al.*, manuscript in revision). In the latter system, a PAM-attenuated SpyCas9 (SpyCas9^MT3^) fused to a programmable DNA-binding domain (pDBD) enables SpyCas9 editing to be restricted to the intended on-target site, with off-target editing suppressed^39^. Zinc-finger proteins, TALE proteins^39^, or orthogonal dCas9s can each serve as the pDBD. AcrIIC4_*Hpa*_ or AcrIIC5_*Smu*_ both suppressed genome editing by SpyCas9^MT3^-dNmeCas9 (Supplementary Fig. 6a), indicating that these type II-C anti-CRISPRs can cross sub-type boundaries and serve as SpyCas9 editing off-switches in this context. As expected, AcrIIC1 orthologs do not inhibit editing by SpyCas9^MT3^-dNmeCas9, either because they act downstream of dNmeCas9 DNA binding^19^, or because they fail to bind to the mutated HNH domain of dNmeCas9 (Supplementary Fig. 6a). SpyCas9 fusion to nuclease-active NmeCas9 can also be used to induce efficient segmental deletion via simultaneous DNA cleavage by both orthologs (Bolukbasi *et al.*, manuscript in revision) (Supplementary Fig. 6b). Type II-C anti-CRISPRs specifically inhibit editing activity of NmeCas9 in the fusion context, leading to the appearance of small indels at the SpyCas9 site rather than a segmental deletion (Supplementary Fig. 6b). These data from mammalian cells confirm the potential utility of AcrIIC4_*Hpa*_ or AcrIIC5_*Smu*_ (as well as other type II anti-CRISPRs) proteins for modulating Cas9-dependent genome engineering applications across subtypes.

## Discussion

The prevalence of CRISPR-Cas immune systems in bacteria and archaea has driven phages to evolve anti-CRISPR proteins. Indeed, numerous anti-CRISPRs against type I and type II systems have been discovered in both bacteria and archaea since the first examples were reported in 2013^15^, with a range of inhibitory mechanisms for impairing Cas protein function^2–4^, ^24^. Here, we report two new families of type II-C anti-CRISPR proteins, AcrIIC4_*Hpa*_ and AcrIIC5_*Smu*_, and their cognate Cas9 proteins from *H. parainfluenzae* and *S. muelleri*. We define PAMs for the newly characterized HpaCas9 and SmuCas9 orthologs and show that they are functional *in vitro* and that HpaCas9 confers phage immunity in bacteria, expanding the functional Cas9 repertoire. These additional anti-CRISPRs and Cas9s total five anti-CRISPR families that differentially inactivate five different type II-C Cas9 orthologs (Fig. 6).

**Figure 6.**
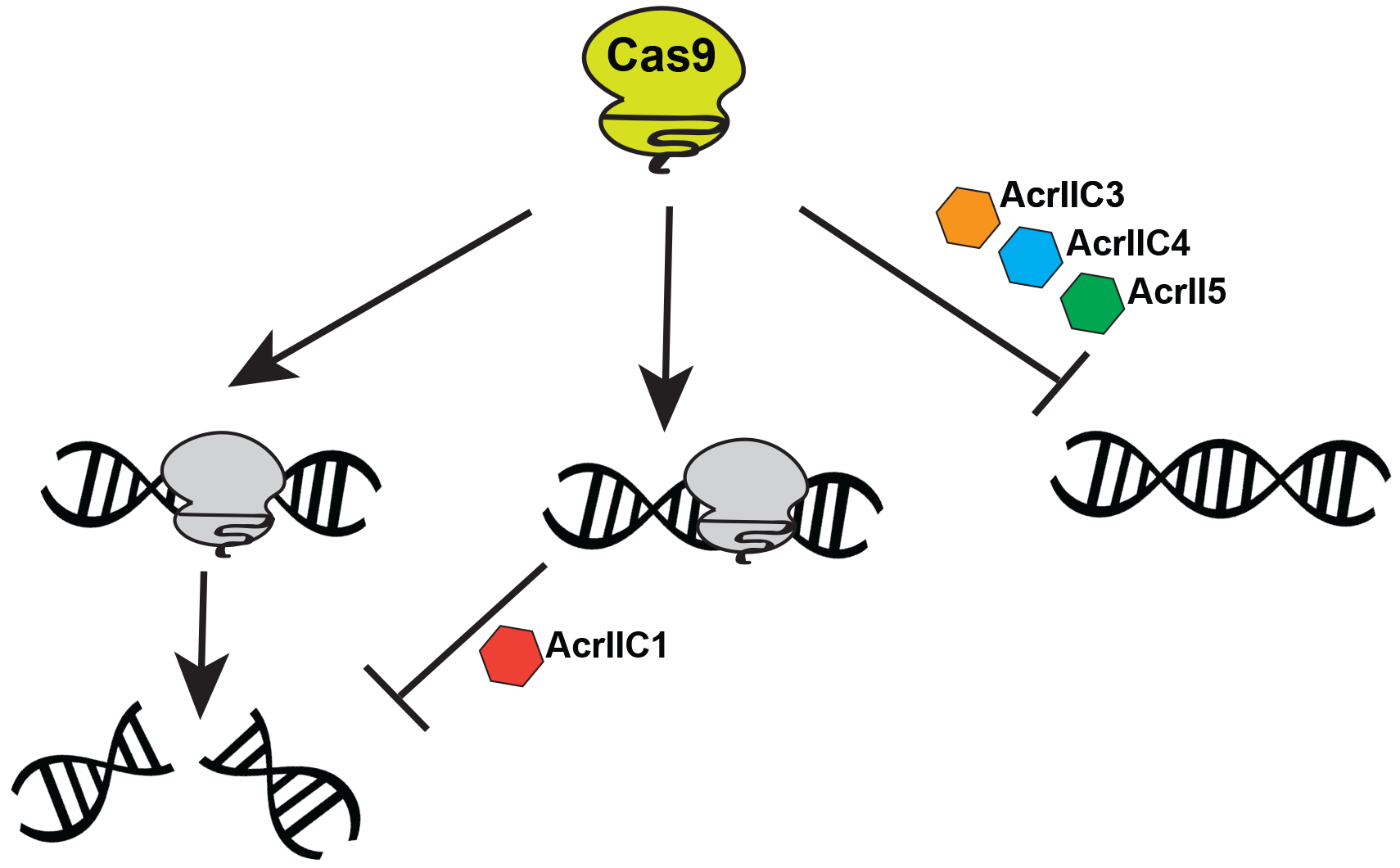
Summary of type II-C Cas9 orthologs and anti-CRISPR families. Type II-C anti-CRISPRs can act at distinct stages of Cas9-mediated target DNA cleavage. While AcrIIC1 binds to the HNH domain and inhibits all a broad spectrum of Cas9 orthologs, AcrIIC4 and AcrIIC5 prevent DNA binding and a have narrower range of inhibition, similar to AcrIIC3.

AcrIIC4_*Hpa*_ and AcrIIC5_*Smu*_ inhibit NmeCas9, HpaCas9, and SmuCas9 activity *in vitro*, as well as CRISPR interference activity in bacteria, and both also prevent NmeCas9-mediated genome editing in mammalian cells. The two new Acr families are the most potent among the type II-C Acrs, prevent substrate DNA binding by NmeCas9 and dNmeCas9, and exhibit higher specificities for inhibition of type II-C Cas9s in comparison to AcrIIC1^19^. AcrIIC4_*Hpa*_ and AcrIIC5_*Smu*_ activity was found to be specific to closely related Cas9 orthologs, as they did not inhibit the more distantly related CjeCas9 and GeoCas9 type II-C orthologs. Cross-species inhibitory effects of each Acr may be graded depending on the similarity of the Cas9 orthologs. Subtle differences may be sufficient to distinguish each anti-CRISPR’s breadth of inhibition as broad-spectrum or highly specific, with gradations between these two extremes. For example, the AcrIIC1 family of Acrs can inhibit multiple Cas9s, likely because they bind to the highly conserved HNH domain^19^, whereas other type II-C Acrs may bind to Cas9 domains that are less conserved (like the PID). The evolutionary pressure on Cas9s to evolve away from anti-CRISPR inhibition may promote diverse PAM specificities and other mechanistic distinctions between Cas9 orthologs. Similarly, distinct anti-CRISPR specificities for inhibiting Cas9 orthologs could suggest different mechanisms of action. We show that AcrIIC4_*Hpa*_ and AcrIIC5_*Smu*_ prevent binding of Cas9 to target DNA, like AcrIIC3 and AcrIIA4 but unlike AcrIIC1^1, 19^. Target DNA binding could be prevented by precluding initial recognition of the PAM (similar to the strategy of AcrIIA4^20–22^), by preventing one of the stages of R-loop formation and Cas9 structural rearrangement^40^, or a combination of these.

Moreover, we have demonstrated the potential utility of Acr-mediated control of Cas9 and dCas9-based technologies by AcrIIC4_*Hpa*_ and AcrIIC5_*Smu*_. Recently, AcrIIA4^17^ was used as an inhibitor of dSpyCas9 fused to a DNA demethylase, Tet1, to inactivate dSpyCas9-Tet1 DNA target binding^41^. Separately, AcrIIA families were shown to prevent a gene-drive propagation in *Saccharomyces cerevisiae*^42^. These are a few examples of the potential utility of Acrs as Cas9 off-switches. Many applications stand to benefit from increasing the numbers, specificities, and inhibitory mechanisms of anti-CRISPRs, for instance through combinatorial control over multiple Cas9/dCas9 proteins. For example, both broad-spectrum (e.g., AcrIIC1_*𝒩me*_) and highly specific (e.g., AcrIIC3_*𝒩me*_, 4_*Hpa*_, or 5_*Smu*_) anti-CRISPR proteins could be used to control multiple Cas9s simultaneously, or specific Cas9s but not others, upstream or downstream of target recognition, to achieve maximal flexibility of both genome manipulation and regulation.

Apart from potential uses in biotechnology, CRISPR-Cas systems and anti-CRISPR proteins that inactivate them are in strong accord with the Red Queen hypothesis, which proposes that bacteria must evolve new mechanisms to resist invaders while the invaders simultaneously evolve countermeasures. The widespread prevalence and extreme diversity of CRISPR-Cas9 systems in bacteria and archaea, as well as the adaptive nature of the resulting defenses, pose a significant challenge to phages and other MGEs. Anti-CRISPR proteins provide phages with an effective tactic to inactivate CRISPR-Cas systems and likely contribute to phage persistence in the face of host defense mechanisms. Many gaps remain in our understanding of the origins of these anti-CRISPRs and how they function in the context of phage predation. It is likely that these proteins have emerged independently and repeatedly through convergent evolution, as indicated by a lack of sequence or structural similarities among many reported Acrs^2–4, 24^. A structural study of a capsid protein from a phage that infects *Thermus thermophilus* shares a common core ß-barrel domain with AcrIIC1, suggesting an evolutionary source for an anti-CRISPR protein^43^. Our ability to address these outstanding questions is limited by the relatively small number of examples of known anti-CRISPR proteins and their striking diversity in sequence and structures. Expanding the collection of diverse anti-CRISPR families and their cognate CRISPR effectors will help further our understanding of the arms race between phages and their hosts.

## Methods

### Characterization of HpaCas9 and SmuCas9

CRISPRfinder (http://crispr.i2bc.paris-saclay.fr) was used to identify the CRISPR locus of *Haemophilus parainfluenzae*. The spacers targeting the phage sequences were blasted via CRISPRTarget (http://bioanalysis.otago.ac.nz/CRISPRTarget) to predict the PAM present on the 3’ sequences. DNA and protein sequences of HpaCas9 and SmuCas9 orthologs are provided in Supplementary Table 1.

### Plasmid Construction

Plasmids used in this study are described in Supplementary Table 4.

#### Cas9/sgRNA and anti-CRISPR vector for bacterial expression, protein purification and in vitro transcription

The pMCSG7-NmeCas9 expression vector and the sgRNA for *in vitro* transcription are as previously described^1^. To make the HpaCas9 expression vector pEJS-MCSG7-HpaCas9, genomic DNA sequence from *H. parainfluenzae* DSM 8987 was obtained from DSMZ and cloned into the pMCSG7-NmeCas9 expression plasmid, replacing the NmeCas9 sequence using Gibson Assembly (NEB). The GeoCas9-expressing plasmid (expressing the GeoCas9 ortholog from *G. stearothermophilus* strain ATCC 7953) was obtained from Addgene (#87700) and similarly cloned into the pMCSG7 vector. To make GeoCas9 from strain L300, a gBlock (IDT) containing the PID was used to replace the PID of GeoCas9 from *G. stearothermophilus* strain ATCC 7953. For construction of sgRNA scaffolds for HpaCas9 and GeoCas9, the tracrRNA was predicted by crRNA repeat complementarity as well as homology to the NmeCas9 tracrRNA. These sgRNA scaffolds were ordered as gBlocks (IDT) along with overhangs to clone into pLKO.1 plasmid^1, 39^ using Gibson Assembly (NEB). The CjeCas9 sgRNA plasmid was used as previously reported^19, 44^. All sgRNA scaffolds were used as templates to create *in vitro*-transcribed sgRNAs.

DNA sequences encoding candidate anti-CRISPR proteins were synthesized and cloned into a pUC57 mini (AmpR) vector with a *𝒩. meningitidis* 8013 Cas9 promoter sequence for bacterial work, as done previously for other anti-CRISPRs^1^. For anti-CRISPR protein purification, the Acr insert was amplified and inserted into the pMCSG7 backbone by Gibson Assembly (NEB), resulting pMCSG7-Acr. Supplementary Table 1 contains the DNA and protein sequences of the anti-CRISPRs tested in this study.

#### Cas9/sgR𝒩A and Acr vectors for mammalian expression

For editing of genomic dual target sites by both SpyCas9 and NmeCas9, we used Cas9 and cognate sgRNA expression vectors that are described previously^1^. To generate the Acr expression vector, the Acr ORF was amplified from pUC57-Acr and inserted into XhoI-digested pCDest2 by Gibson assembly (NEB).

#### Vectors for fluorescence microscopy

pHAGE-TO-DEST dSpyCas9-(mCherry)_3_ and dNmeCas9-(sfGFP)_3_ plasmids^38^ were purchased from Addgene (#64108 and #64109, respectively) and used directly for no-sgRNA control experiments. dNmeCas9-(sfGFP)_3_ and dSpyCas9-(mCherry)_3_ all-in-one plasmids have been described previously (Pawluk et al., 2016). To make Acr plasmids, we amplified an mTagBFP2 cassette and incorporated it into pCDest2 vectors expressing the respective Acr by Gibson Assembly (NEB).

### Expression and Purification of Acr and Cas9 proteins

The expression and purification of Acrs and Cas9s was performed as described^1, 12^. 6xHis-tagged anti-CRISPRs and Cas9s were expressed in *E. coli* strain BL21 Rosetta (DE3). Cells were grown in LB or 2X YT medium at 37 °C to an optical density (OD_600 nm_) of 0.6 in a shaking incubator. At this stage the bacterial cultures were cooled to 18 °C, and protein expression was induced by adding 1 mM IPTG. Bacterial cultures were grown overnight at 18 °C (~16 hrs), after which cells were harvested and resuspended in Lysis buffer [50 mM Tris-HCl (pH 7.5), 500 mM NaCl, 5 mM imidazole, 1 mM DTT] supplemented with 1 mg/mL Lysozyme and protease inhibitor cocktail (Sigma). Cells were lysed by sonication and the supernatant was then clarified by centrifugation at 18000 rpm for 30 min. The supernatant was incubated with pre-equilibrated Ni-NTA agarose (Qiagen) for 1 hr. The resin was then washed twice with Wash buffer [50 mM Tris-HCl (pH 7.5), 500 mM NaCl, 25 mM imidazole, 1 mM DTT]. The proteins were eluted in Elution buffer containing 300 mM imidazole. For Acr proteins, the 6xHis tag was removed by incubation with His-tagged Tobacco Etch Virus (TEV) protease overnight at 4 °C followed by a second round of Ni-NTA purification to isolate successfully cleaved, untagged anti-CRISPRs (by collecting the unbound fraction). Cas9s were further purified using cation exchange chromatography using a Sepharose HiTrap column (GE Life Sciences). Size exclusion chromatography was used to purify NmeCas9 further in 20 mM HEPES-KOH (pH 7.5), 300 mM KCl and 1 mM TCEP.

### *In vitro* DNA cleavage

For the *in vitro* DNA cleavage experiments with NmeCas9 (Fig. 1C and Supplementary Fig. 1), NmeCas9 sgRNA targeting N-TS4B was generated by *in vitro* T7 transcription (NEB). NmeCas9 (150 nM) was incubated with purified, recombinant anti-CRISPR protein (0-5 μM) in cleavage buffer [20 mM HEPES-KOH (pH 7.5), 150 mM KCl, 1 mM DTT] for 10 minutes. Next, sgRNA (1:1, 150 nM) was added and the mixture was incubated for another 15 minutes. Plasmid containing the target protospacer NTS4B was linearized by ScaI digestion. Linearized plasmid was added to the Cas9/sgRNA complex at 3 nM final concentration. The reactions were incubated at 37 °C for 60 minutes, treated with proteinase K at 50 °C for 10 minutes, and visualized after electrophoresis in a 1% agarose/1xTAE gel.

### Phage immunity

Plasmids expressing Cas9 targeting *E. coli* phage Mu were co-transformed into *E. coli* strain BB101with plasmids expressing the anti-CRISPRs^19^. Cells carrying both plasmids were grown in lysogeny broth (LB) supplemented with streptomycin and chloramphenicol. Anti-CRISPR gene expression was induced using 0.01 mM IPTG for three hours. A lawn of 200 μL of cells in top agar was applied to LB agar plates supplemented with streptomycin, chloramphenicol, 200 ng/mL anhydrotetracycline (aTc), 0.2% arabinose +/− 200 ng/mL aTc, and 10 mM MgSO_4_. Ten-fold serial dilutions of phage Mu were spotted on top of the lawn and the plates were incubated overnight at 37 °C. To confirm the expression levels of the anti-CRISPR proteins in this assay, 500 μL aliquots of cells applied to the top agar were pelleted by centrifugation, resuspended in 100 μL of SDS-PAGE loading buffer, run on a 15% Tris-Tricine gel, and the resulting protein gel was visualized by Coomassie Blue.

### Cas9-Acr co-purification

Cas9 proteins were expressed from plasmid pMCSG7 with an N-terminal 6-His affinity tag in *E. coli* Rosetta cells. Untagged Acrs were co-expressed in the same cells from plasmid pCDF1b. Cells were grown in LB to an OD_600_ of 0.8 and protein production was induced with 2 mM IPTG overnight at 16 °C. Cells were collected by centrifugation, resuspended in binding buffer (20 mM Tris, pH 7.5, 250 mM NaCl, 5 mM imidazole), lysed by sonication, and cellular debris was removed by centrifugation. The cleared lysates were applied to Ni-NTA columns, washed with binding buffer supplemented with 30 mM imidazole, and eluted with 300 mM imidazole. Protein complexes were analyzed by SDS-PAGE followed by Coomassie staining.

### PAM discovery

A library of a protospacer with randomized PAM sequences was generated using overlapping PCRs, with the forward primer containing the 10-nt randomized sequence flanking the protospacer. The library was subjected to *in vitro* cleavage by purified recombinant HpaCas9 or SmuCas9 proteins as well as *in vitro* transcribed sgRNAs. Briefly, 300 nM Cas9:sgRNA complex was used to cleave 300 nM of the target fragment in 1X reaction buffer (NEBuffer 3.1) at 37 °C for 60 minutes. The reaction was then treated with proteinase K at 50 °C for 10 minutes and run on a 4% agarose gel with 1X TAE. The segment of a gel where the cleavage products were expected to be was purified and subjected to library preparation as described^45^. The library was sequenced using the Illumina NextSeq500 sequencing platform and analyzed with custom scripts.

### Electrophoretic Mobility Shift Assay (EMSA)

1 μM NmeCas9 was incubated with 1 μM sgRNA in 1x Binding buffer [20 mM Tris-HCl (pH 7.5), 150 mM KCl, 2 mM EDTA, 1 mM DTT, 5% glycerol, 50 μg/mL Heparin, 0.01% Tween 20, 100 ug/mL BSA] for 20 minutes at room temperature to form the RNP complex. Acrs were added to a final concentration of 10 μM and incubated for an additional 20 minutes. Finally the FAM-tagged NTS4B protospacer oligonucleotide was added to the mixture and incubated at 37 °C for 1 hr. The mixture was loaded onto a native 6 % acrylamide gel and the FAM-tagged DNA was visualized using a Typhoon imager.

### sgRNA EMSA

NmeCas9 (1.5 μM) and anti-CRISPR (20 μM) proteins were pre-incubated in 1x Binding buffer for 10 minutes and then sgRNA (0.15 μM) was added to the reaction for additional 10 minutes. The complexes were resolved on a 6% polyacrylamide native gel, stained by SYBR Gold (ThermoFisher) and visualized with a Typhoon imager.

### Mammalian genome editing

Plasmids for mammalian expression of NmeCas9, SpyCas9, their respective sgRNAs, and the anti-CRISPR proteins are listed in Supplementary Table 4. Plasmid transfections, collection of genomic DNA, and T7E1 digestions were as described^1^.

### Genome editing by Cas9 ribonucleoprotein (RNP) delivery

RNP delivery of NmeCas9 was performed using a Neon electroporation system following the manufacturer’s instructions (ThermoFisher). Briefly, in a 10 uL reaction volume, 15 pmol of NmeCas9 and 150 pmol of anti-CRISPR protein were mixed in buffer R and incubated at room temperature for 20 minutes. 20 pmol of T7 *in vitro*-transcribed sgRNA was added to the Cas9-Acr complex and incubated at room temperature for 30 minutes. Approximately 50,000-100,000 cells were mixed with the RNP-Acr-sgRNA complex, electroporated (Neon nucleofection system), and then plated in 24-well plates. Genomic DNA was extracted 48 hours post-nucleofection using a DNeasy Blood and Tissue kit (Qiagen) according to the manufacturer’s protocol. Quantification of editing (% of amplicons exhibiting lesions) was done using TIDE analysis35. PCR products spanning the target site were amplified using 2x HiFi master mix (NEB) and column-purified (Zymo). Purified amplicons were sent for Sanger sequencing (Genewiz) and trace files were analyzed by TIDE.

### Fluorescence microscopy of dNmeCas9

Experimental procedures were as described^1^. Briefly, U2OS cells were co-transfected with all-in-one plasmids (150 ng of each dNmeCas9 and dSpyCas9 plasmid), additional sgRNA-expressing plasmid, and 100ng of anti-CRISPR/mTagBFP2 plasmid using PolyFect (Qiagen) according to the manufacturer’s instructions. After 24 hours of incubation, live cells were imaged with a Leica DMi8 microscope equipped with a Hamamatsu camera (C11440-22CU), a 63x oil objective lens, and Microsystems software (LASX). Further imaging processing was done with Fiji-ImageJ. For quantification, only cells that exhibited mTagBFP2 and sfGFP fluorescence as well as dSpyCas9-(mCherry)_3_ telomeric foci were assessed for the presence or absence of co-localizing dNmeCas9-(sfGFP)_3_ telomeric foci.

## Data availability

Raw data files are available upon reasonable request. High-throughput sequencing data is in preparation for deposition in the NCBI Sequence Read Archive (SRA).

## Acknowledgements

We are grateful to Y. Hidalgo-Reyes for technical assistance, and to members of the Davidson, Maxwell and Sontheimer labs for helpful discussions. We also thank M.F. Bolukbasi and S.A. Wolfe for sharing unpublished data and resources. This work was supported by grants from the Canadian Institutes for Health Research to A.R.D. (FDN-15427) and K.L.M. (PJT-152918), and by a grant from the U.S. National Institutes of Health (GM125797) to A.R.D. and E.J.S.

## Author contributions

J.L. carried out co-immunoprecipitations, western analyses, and fluorescence microscopy experiments. A.E. and A.M. characterized *H. parainfluenzae* and *S. muelleri* CRISPR loci and expressed, purified, and analyzed HpaCas9 and SmuCas9. A.E. and I.G. designed and executed PAM definitions. A.M., J.L. and H.E.L. expressed and purified anti-CRISPR and NmeCas9 proteins, and A.M. and H.E.L. conducted *in vitro* analyses of Acr proteins. B.G. designed, performed, and analyzed phage and co-purification binding assays. J.L., N.A., R.I. and X.D.G. designed, performed, and analyzed mammalian genome editing experiments, and P.L. analyzed targeted deep sequencing data. A.P. performed bioinformatic analyses identifying candidate anti-CRISPRs. A.R.D., K.L.M., and E.J.S. supervised experiments. J.L., K.L.M. and E.J.S. wrote the manuscript, and all authors edited the manuscript.

